# Effect of EBV-Transformation on Oxidative Phosphorylation Physiology in Human Cell lines

**DOI:** 10.1101/2019.12.16.878025

**Authors:** Rachel N. Weinstein, Douglas L. Crawford

## Abstract

Do the immortalized and cryopreserved white blood cells that are part of the 1,000 Human Genomes Project represent a valuable cellular physiological resource to investigate the importance of genome wide sequence variation? While much research exists on the nucleotide variation in the 1,000 Human Genomes, there are few quantitative measures of these humans’ physiologies. Fortunately, physiological measures can be done on the immortalized and preserved cells from each of the more than 1,000 individuals that are part of Human Genome project. However, these human white blood cells were immortalized by transforming them with the Epstein-Barr virus (EBV-transformed lymphoblastoid cell lines (LCL)). This transformation integrates the viral genome into the human genome and potentially affects important biological differences among individuals. The questions we address here are whether EBV transformations significantly alters the cellular physiology so that 1) replicate transformations within an individual are significantly different, and 2) whether the variance among replicates obscures the variation among individuals. To address these questions, we quantified oxidative phosphorylation (OxPhos) metabolism in LCLs from six individuals with 4 separate and independent EBV-transformations. We examined OxPhos because it is critical for energy production, and mutations in this pathway are responsible for most inborn metabolic diseases. The data presented here demonstrate that there are small but significant effects of EBV-transformations on some OxPhos parameters. In spite of significant variation due to transformations, there is greater and significant variation among individuals in their OxPhos metabolism. Thus, the LCLs from the 1,000 Human Genome project could provide valuable insights into the natural variation of cellular physiology because there is statistically significant variation among individuals when using these EBV-transformed cells.

## Introduction

Heritable metabolic disease affecting mitochondrial respiration and the oxidative phosphorylation (OxPhos) pathway occur at a high frequency: 1/5000 individuals [1–7]. Many mitochondrial diseases are caused by point mutations in one of the 13 mitochondrial encoded proteins or mutations in the mt-tRNAs [4, 8, 9]. Most of these mtDNA mutations have an adult disease onset and can have very specific cellular targets (*e.g.*, lesions in a specific area of the basal ganglion [4, 8, 9]). Along with the 13 mtDNA genes, 76 nuclear genes encode proteins for the 5 OxPhos enzyme complexes (Figure 1). In contrast to mtDNA mutations, deleterious mutations in these nuclear genes affect more individuals, have greater penetrance, are typically lethal, and have early onset (within the first year of life; [4–6]). Surprisingly, mtDNA mutations affecting OxPhos proteins have a heterogeneous disease presentation and variable penetrance even among homoplasmic individuals [2, 4, 5], and many individuals have nuclear SNPs associated with pathogenic diseases but remain healthy [10–13]. These observations suggest that interactions among DNA polymorphisms in both mitochondrial and nuclear genomes have significant effects on human health by affecting the OxPhos pathway. To better understand these observations would involve quantifying the natural variation in OxPhos metabolism taking into consideration genomic polymorphisms.

**Figure 1.**
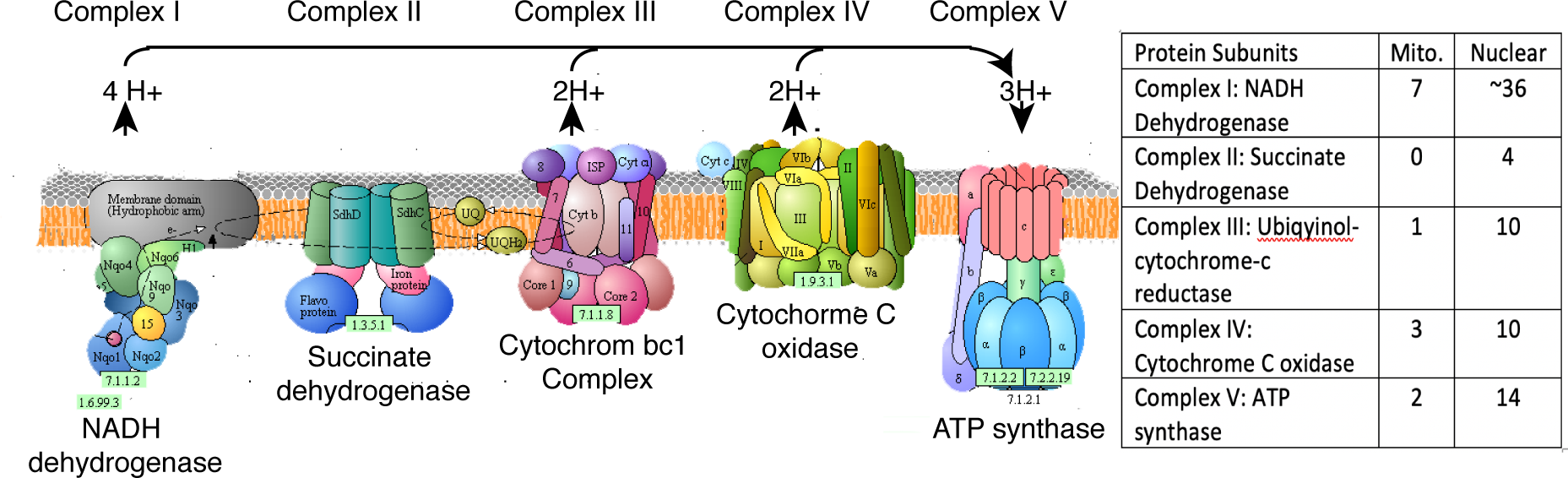
OxPhos pathway. OxPhos consists of five enzyme complexes with both mitochondrial and nuclear genomes encode proteins in four of these complexes. Numbers of mitochondrial and nuclear encoded proteins in the different subunits are listed. http://www.genome.jp/kegg-bin/show_pathway?map00190 [31–33]

Quantitative analyses of OxPhos metabolism with knowledge about DNA sequence variation in OxPhos genes and the approximately 1,5000 other nuclear genes that affect mitochondrial ATP production [1] is possible using the resources from the “1,000 Human Genome” project [14]. The individuals in the 1,000 Human Genome collection represent much of the genetic diversity in humans worldwide, with sampled populations ranging from East and South Asia, Europe, West Africa, and the Americas [14]. Importantly, in addition to having fully sequenced genomes, these individuals have cryopreserved, immortalized, lymphoblastoid cell lines (LCL). These LCLs originate as peripheral B lymphocytes that have been isolated from human subjects and transformed with the Epstein-Barr virus (EBV) [15].

Transformed LCLs are effectively immortal, express a wide array of genes including many metabolic pathways, and are easy to culture in a laboratory setting [15]. EBV-transformed LCLs and specifically cell lines from the 1,000 Human genome project have been used successfully to define sex bias in gene expression [16], statin-dependent QTL [17], cell specific networks [18], genetic variation in DNA replication [19] and epigenetics [20, 21]. Overall, these and other studies have concluded that EBV-transformed LCLs continue to represent naturally occurring biological variation [22, 23]. However, EBV-transformation can have significant effects on gene expression, and there is considerable debate on whether this transformation alters biologically important phenotypic differences, including cellular physiology, which would reduce the EBV-transformed LCLs’ utility for understanding the effects of nucleotide variation [20, 24, 25]. Because it is unclear if the random viral integration affects physiological variation in LCLs, we sought to quantify the variation that arises from this transformation by quantifying the metabolic activity of the oxidative phosphorylation (OxPhos) pathway. This approach would have been strengthened by comparing pre-transformed with the currently available transformed LCLs, but these untransformed cells are unavailable. Instead, presented here we investigated whether independently transformed LCLs within individuals introduces significantly large amount of variance such that there is little remaining variation among individuals (i.e. variation within individual is greater than the variation among individuals).

To investigate the effect of EBV-transformation, we used six individual donors each with four, independent, replicate EBV-transformed cell lines [26]. These independent EBV-transformations are used to quantify the effect of transformation and whether EBV-transformation obscured significant variation among individuals. For our study we focused on OxPhos metabolism, and we measured the six parameters of the OxPhos pathway: State 3 activity, E State activity, Complex I activity, Complex II activity, Complex IV activity, and proton leak across the mitochondrial membrane. Overall, EBV-transformation had a small but significant effect, yet this effect did not obscure differences among individuals.

### Methods and Materials

### Cell Culturing

Table 1 lists the six individuals each with four independent LCLs. EBV-transformed LCLs were obtained from NIGMS Human Genetic Cell Repository at the Coriell Institute for Medical Research [26] [27]. Six different individual’s cell lines were grown and assayed weekly followed by the same individual with different transformations in subsequent weeks. For each cell line, typically three replicates were measured over a five-day period. Thus, there are three replicates per cell line, four cell lines per individual, and six individuals.

**Table 1.**
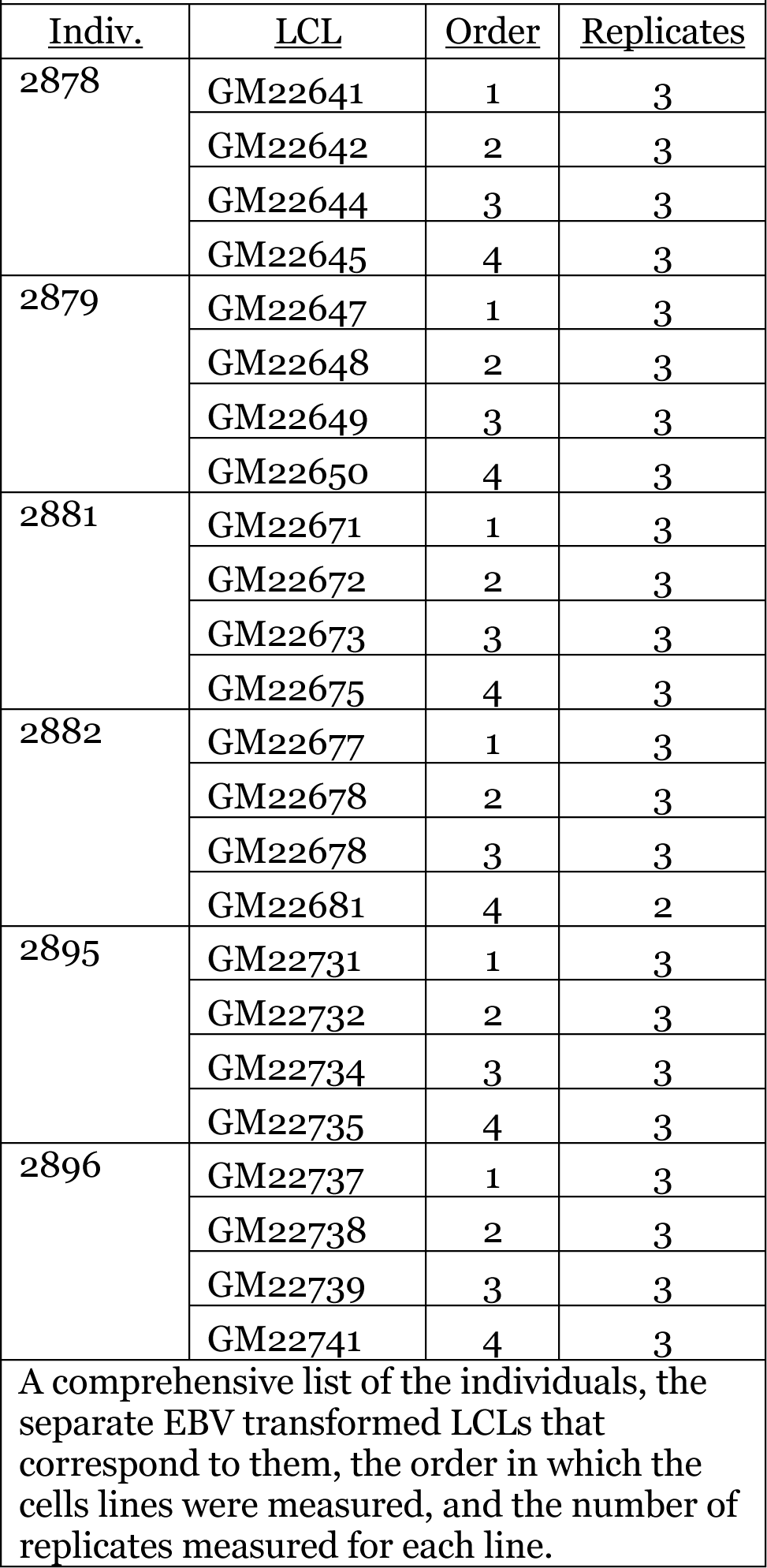
Individuals and Lymphoblastoid Cell Lines

| <u>Indiv.</u> | <u>LCL</u> | <u>Order</u> | <u>Replicates</u> |
| --- | --- | --- | --- |
| 2878 | GM22641 | 1 | 3 |
|  | GM22642 | 2 | 3 |
|  | GM22644 | 3 | 3 |
|  | GM22645 | 4 | 3 |
| 2879 | GM22647 | 1 | 3 |
|  | GM22648 | 2 | 3 |
|  | GM22649 | 3 | 3 |
|  | GM22650 | 4 | 3 |
| 2881 | GM22671 | 1 | 3 |
|  | GM22672 | 2 | 3 |
|  | GM22673 | 3 | 3 |
|  | GM22675 | 4 | 3 |
| 2882 | GM22677 | 1 | 3 |
|  | GM22678 | 2 | 3 |
|  | GM22678 | 3 | 3 |
|  | GM22681 | 4 | 2 |
| 2895 | GM22731 | 1 | 3 |
|  | GM22732 | 2 | 3 |
|  | GM22734 | 3 | 3 |
|  | GM22735 | 4 | 3 |
| 2896 | GM22737 | 1 | 3 |
|  | GM22738 | 2 | 3 |
|  | GM22739 | 3 | 3 |
|  | GM22741 | 4 | 3 |

A comprehensive list of the individuals, the separate EBV transformed LCLs that correspond to them, the order in which the cells lines were measured, and the number of replicates measured for each line.

The cells were cultured in 18 mL of RPMI 1640 media (with 2mM L-glutamine and 15% fetal bovine serum) in upright T25 tissue culture flasks and kept at 37°C, 70% humidity, and 5% carbon dioxide. Cell lines were cultured for 2 weeks before OxPhos measurements to ensure that they had sufficient acclimation in the provided media. Cells were diluted approximately every 3 days between 30-70% to achieve 300,000 cells/mL. The day prior to OxPhos determination cells were diluted to 500,000 cells/mL, and the cells would grow to approximately 1,000,000 cells/mL on the day of measurement. 1.5 million cells were used to measure OxPhos metabolism.

The cell concentration in a given culture was estimated by staining a sample with a 2:1 ratio of cells to Trypan blue dye for 3-5 minutes. The cells were then counted on a glass haemocytometer. DNA concentration was determined and used as the measure of cell density for each LCL sample used for OxPhos measurements.

### Oxidative Phosphorylation Measurements

The OxPhos metabolic parameters are defined in Table 2. For OxPhos measurements, approximately 3,000,000 cells were spun down at 200 x G for 12 minutes and brought up at 1.5 x10^7^ cell/mL in MiR05 (3mM MgCl2, 60mM K-lactobionate, 0.5 mM EGTA, 20 mM Taurine, 10mM KH2PO4, 20mM Hepes pH, 7.1 @22°C, 110mM sucrose, 1g/liter fatty acid free BSA). The cells’ walls were then permeated with 7.5 μL of 10 mg/mL digitonin (for 10 minutes at 37°C) to allow reagents access to the cells’ mitochondria. One hundred microliters (1.5 x10^6^ cells) of these permeated cells were used. Six parameters of the OxPhos pathway were measured (Table 2): State 3 activity, E State activity, Complex I activity, Complex II activity, Complex IV activity, and proton leak across the mitochondrial membrane. A seventh parameter, QC, measures the effect of added cytochrome c, which for intact mitochondria should have negligible effect. For this study the ratio of added cytochrome C/State 3 was always 1 (+/− 5%), and thus the cell preparations maintained mitochondrial intactness. These parameters were measured via oxygen consumption using the Oxygraph-2k and Datlab software (Oroboros Instruments, Innsbruck, Austria). For each measurement, 1.5 x10^6^ cells were added to a chamber in the Oxygraph-2k containing 2mL of Miro5 at 37°C. First, to determine State 3 activity, substrates and ADP were added for a final concentration of 5mM pyruvate, 10mM glutamate, 10 mM of succinate, and 2.5mM of ADP (adenosine 5’ diphosphate). Second, cytochrome c (1ouM final) was added to assess mitochondrial membrane integrity. Third, oligomycin (2uM final) – a Complex V, or ATP synthase inhibitor - was added to determine proton leak across the mitochondrial membrane. Fourth, FCCP (carbonyl cyanide-4-phenylhydrazone; 0.025uM final) – a mobile ion carrier – was added to determine E State., Fifth, 1 μL of rotenone (0.05uM final) - a Complex I inhibitor - was added, followed by malonic acid (5mM) - a Complex II inhibitor - to determine the activity of Complex II., Sixth, 1uL of antimycin A (2.5uM final) – a Complex III inhibitor - was added. Finally, 10 μL of 1:1 TMPD (N, N, N’, N’-tetramethyl-p-phenylenediaminedihydrochloride; 0.5 mM final) and ascorbate (2mM final) were added, which determines Complex IV activity. Three replicate measures on different days were determined for each transformation from each individual.

**Table 2.**
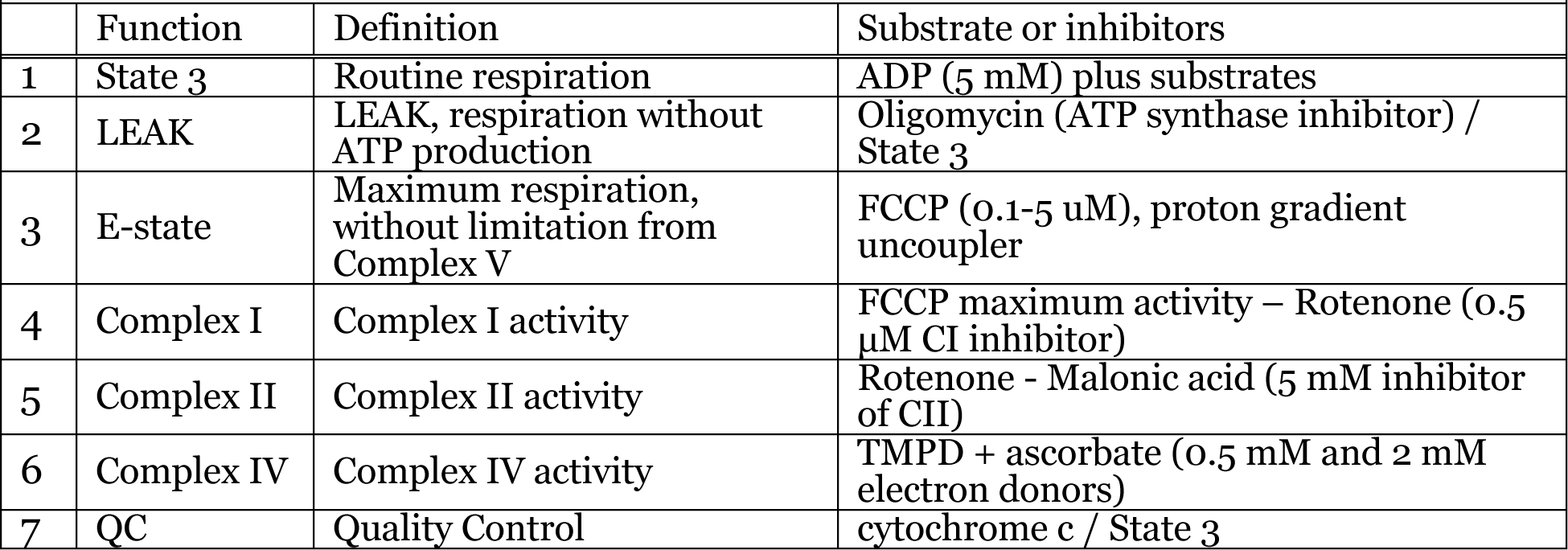
OxPhos Functions

| **Table 2.** OxPhos Functions |  |  |  |
| --- | --- | --- | --- |
|  | Function | Definition | Substrate or inhibitors |
| 1 | State 3 | Routine respiration | ADP (5 mM) plus substrates |
| 2 | LEAK | LEAK, respiration without ATP production | Oligomycin (ATP synthase inhibitor) / State 3 |
| 3 | E-state | Maximum respiration, without limitation from Complex V | FCCP (0.1-5 uM), proton gradient uncoupler |
| 4 | Complex I | Complex I activity | FCCP maximum activity – Rotenone (0.5 µM CI inhibitor) |
| 5 | Complex II | Complex II activity | Rotenone - Malonic acid (5 mM inhibitor of CII) |
| 6 | Complex IV | Complex IV activity | TMPD + ascorbate (0.5 mM and 2 mM electron donors) |
| 7 | QC | Quality Control | cytochrome c / State 3 |

### DNA Concentration Measurements

DNA concentration was used as a more accurate measurement of the number of cells measured in the Oxygraph-2k. For this assay, 50 μL of cells used for OxPhos determinations were stored at −80°C in 100 μL of 2% SDS-TE prior to determination of DNA concentration. The DNA concentration assay was performed using the Biotium AccuBlue High Sensitivity DNA kit.

### Statistical Analyses

The normal distribution of all six OxPhos parameters (Table 2) and DNA concentration were tested using “Goodness Fit” (JMP, SAS) with the null hypothesis that data fit a normal distribution. Normal distribution was rejected for DNA concentration and 5 of 6 OxPhos parameters: State 3, E State, Complex I, Complex II, and proton LEAK. Log_10_ transformation of these parameters were normally distributed (p >0.01 Goodness of Fit). Thus, data analyses used log_10_ transformation. Unlike State 3, E State, Complex I, Complex II, and proton LEAK data, Complex IV data were normally distributed using the original values, so untransformed values were used. Additionally, simply dividing metabolic activity by DNA concentration did not remove the effect of DNA: i.e., DNA had significant covariance with the OxPhos parameter when divided by DNA. Dividing the activity of our OxPhos parameters by logarithmic value of the DNA concentration (Log[DNA]) removed this significance—that is, the OxPhos parameter divided by Log[DNA] relative to [DNA] or Log[DNA] had zero slope. Thus, all OxPhos parameters (both log and non-log transformed) are expressed as a ratio with Log[DNA] as the denominator.

For each OxPhos parameter, a nested ANOVA analysis was used to quantify differences between independent transformations within an individual and differences between individuals. Thus, for the nested ANOVA three replicates per cell line were nested within the four cell lines per individual, and these four independent EBV-transformations were nested within the six individuals. Differences among individuals for each parameter were further examined using a Tukey post-hoc test. Furthermore, a one-way ANOVA analysis was performed within each individual to identify the EBV-transformed specific differences within an individual. All analyses used JMP software (SAS Cary, NC).

## Results

A total of 24 cells lines were analyzed, consisting of 6 individuals each with 4 independent EBV-transformations. Each LCL had OxPhos activity measured in triplicate, with the exception of one cell line (individual 2882, transformation GM22681) that was measured in duplicate (Table 1).

Table 3 provides nested ANOVA results (individuals, transformation nested within individuals) for all 6 OxPhos parameters. State 3 respiration (overall mitochondrial respiration through the OxPhos pathway) showed significant differences among EBV-transformations within an individual (p=0.0006) and among individuals (p=0.0078; Table 4A). Similarly, E State (maximal OxPhos respiration), Complex I, and Complex IV dependent respirations showed significant variation among EBV-transformations within an individual and among individuals (p < 0.02 and p < 0.05, respectively; Table 3 B, C and E). Complex II however, only showed significant variation in activity between transformations within an individual (p=0.0061) and not between individuals (p=0.0705, Table 3E). Proton LEAK across the mitochondrial membrane, interestingly, appeared to have no significant variation between either separate transformations within an individual (p=0.5634) or among individuals (p=0.7074; Tables 3F).

**Table 3.** Nested ANOVA results of OxPhos parameters (transformation within individuals and individuals). The degrees of freedom, sum of squares, F ratios, and p values determined for each measured OxPhos parameter in both individuals and separate transformations within individuals.

| *A: State 3 Metabolism* |  |  |  |  |
| --- | --- | --- | --- | --- |
|  | *df* | *ss* | *F* | *p* |
| Individuals | 5 | 0.0328 | 3.5932 | **0.0078\*\*** |
| Transformations | 18 | 0.1064 | 3.2343 | **0.0006\*\*** |
| *B: E State Metabolism* |  |  |  |  |
|  | *df* | *ss* | *F* | *p* |
| Individuals | 5 | 0.0281 | 2.7142 | 0.0310* |
| Transformations | 18 | 0.1069 | 2.8613 | **0.0020\*\*** |
| *C: Complex I Metabolism* |  |  |  |  |
|  | *df* | *ss* | *F* | *p* |
| Individuals | 5 | 0.2171 | 3.7969 | **0.0057\*\*** |
| Transformations | 18 | 0.4562 | 2.2157 | 0.0150* |
| *D: Complex II Metabolism* |  |  |  |  |
|  | *df* | *ss* | *F* | *p* |
| Individuals | 5 | 0.0236 | 2.1957 | 0.0705 |
| Transformations | 18 | 0.0968 | 2.5022 | **0.0061\*\*** |
| *E: Complex IV Metabolism* |  |  |  |  |
|  | *df* | *ss* | *F* | *p* |
| Individuals | 5 | 762.98 | 4.6366 | **0.0016\*\*** |
| Transformations | 18 | 2494.39 | 4.2106 | **<0.0001\*\*\*** |
| Notes. *$p < 0.05$. **$p < 0.01$. ***$p < 0.001$. | | | | |
| *F: Proton Leak* |  |  |  |  |
|  | *df* | *ss* | *F* | *p* |
| Individuals | 5 | 0.0084 | 0.5902 | 0.7074 |
| Transformations | 18 | 0.0473 | 0.9168 | 0.5634 |
| Notes. *$p < 0.05$. **$p < 0.01$. ***$p < 0.001$. **Bold** fonts signify $p < 0.05$ after Bonferroni's multiple correction test across all six parameters | | | | |

**Table 4.** EBV Significant Effect. The three individuals among the six with significant EBV effects. Numbers refer to the order and transformation that was significant (see Table 2).

|  | Individuals with significant EBV effects. |  |  |
| --- | --- | --- | --- |
| OxPhos Parameters | 2879 | 2881 | 2995 |
| State 3 | ns | **2, 3>1** | **1 >3, 4 and 2>4** |
| E-State | ns | ns | **1 >3, 4 and 2>4** |
| Complex _I | ns | ns | **1 >4** |
| Complex _II | ns | ns | **1,2,3 >4** |
| Complex _IV | **2>1** | **2,3,4 >1** | ns |
| LEAK | ns | ns | ns |

Four of the six OxPhos parameters that were significant among individuals (State 3, E State, Complex I and Complex IV, Figure 2). For State 3, E State, and Complex I, individual 2881 typically has the lowest values and 2895 has the highest values (Figure 2). For Complex IV dependent metabolisms, individual 2878 is significantly greater than individuals 2879 and 2881 (Tukey post-hoc test, p < 0.05). To determine which EBV-transformations were significantly different within each individual, we performed a one-way ANOVA for each individual contrasting the four EBV-transformations. Transformation is only significant in 3 individuals: 2879, 2881, and 2895 (Table 4). The other 3 individuals, 2878, 2882, and 2896, did not have a significant effect from transformation. The OxPhos parameters affected by EBV-transformation were State 3 in two of these three individuals (2881 and 2995), E-State, Complex I, and II in individual 2995 and Complex IV in two individuals (2879 and 2881, Table 4). Among these three individuals, the significant effects of transformation were from OxPhos measured in different weeks. That is, although all six individuals were measured in triplicate weekly, the separate transformations were measured in different weeks (Table 2), and order of significance were different for different individuals (Table 4). Individual 2879 showed a significant transformation effect only for Complex IV activity (p=0.0348) where the first assay was significantly less than the second assay (Tukey post-hoc test, p < 0.05). Individual 2881 had significant effects of transformation in State 3 activity (p=0.0154) and Complex IV (p=0.0047), where the first assay is less than the second and third assays (Tukey post-hoc test, p < 0.05). Individual 2895 had significant transformation effects for State 3 activity (p=0.0008), E State, (p=0.0007), Complex I (p=0.0244), and Complex II (F=16.7706; p=0.0008), where the first assay is the highest and the fourth assay is the lowest.

**Figure 2.**
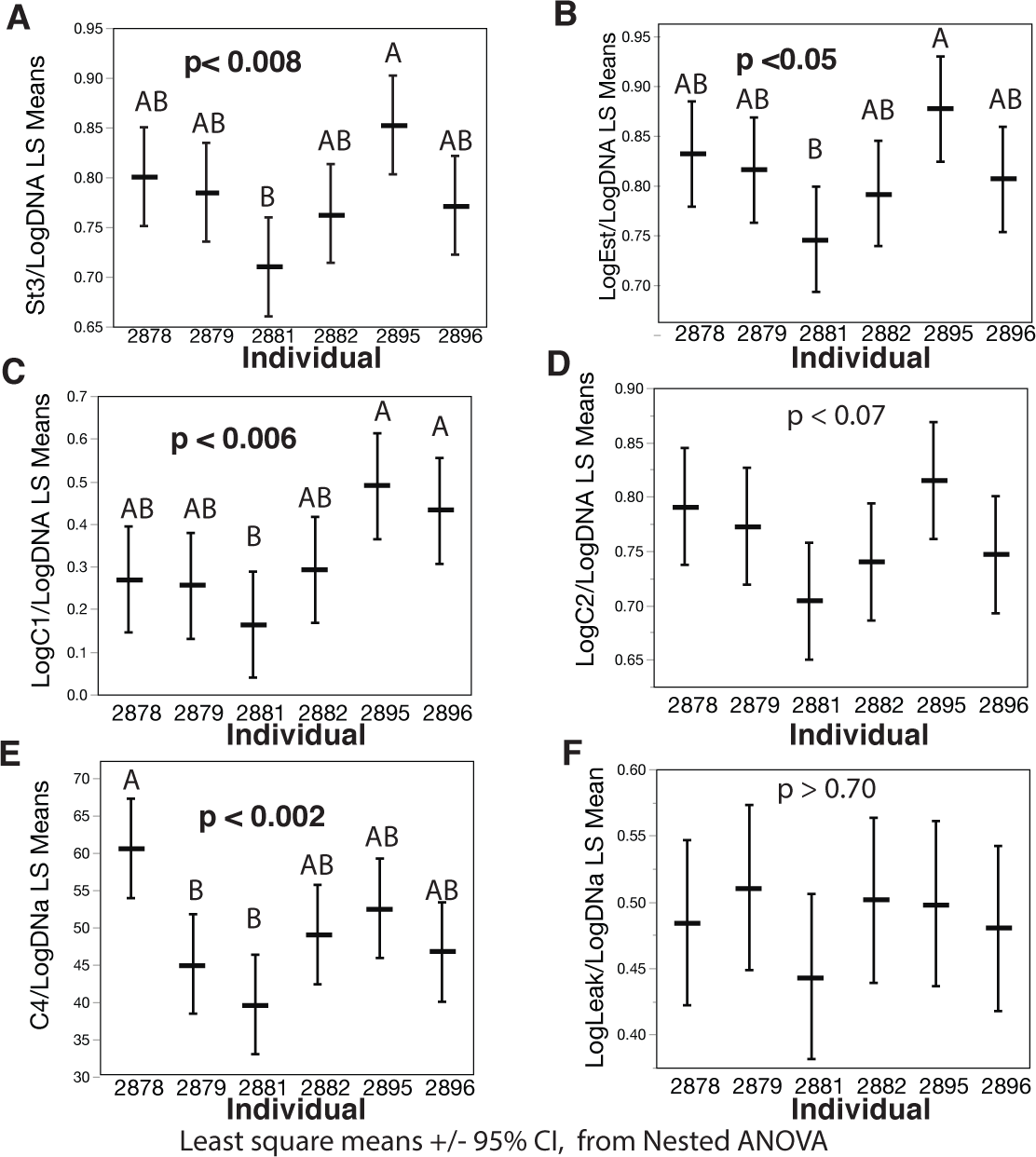
Variation among individuals in OxPhos metabolism. Least-square means and 95% confidence intervals are plotted for the six individuals. “p is the probability of rejecting the null hypothesis of no difference among individuals. Letters above indicate significant groups: different letters are significant (p <0.05, Tukey post-hoc test).

## Discussion

EBV-transformation has had varied effects on LCLs. In the one of the first studies [20], EBV-transformation relative to whole blood had small but significant effects on mRNA expression and methylation patterns, yet LCLs recapitulate the naturally occurring gene expression variation in primary B cells [20]. Similarly in EBV-transformed LCLs cells, cell cycle genes have significantly greater expression compared to primary tissues, which is thought to be the result of less repressive transcription factor regulation [18]. These differences paired LCLs with whole blood and found that the regulatory changes were subtle; for example the levels of mRNA expression for the transcription factor Sp1 and Sp3 were not significantly different in transformed *versus* native cells, but the targets for these transcription factors differed [18]. Furthermore, mRNA expression substantially changed with passage [16, 18, 25]. In contrast, EBV-transformation had no significant effect on mitochondrial copy number, with nearly equal numbers of mitochondria among replicate EBV-transformations [20]. Additionally, EBV-transformation did not obscure sex differences among 99 individuals [16]. Similarly, EBV-transformation did not obscure the association of 16 regulator genes with DNA replications among 161 individual from the EBV-transformed LCLs in 1,000 Human Genome collection [19]. Furthermore, EBV-transform cells are used regularly in pharmacological studies with great success [17, 23]. These data, while acknowledging EBV-transformation effects, find that transformation effects do not inhibit genetics studies. What is unclear is whether these EBV-transformation effects on molecular and biochemical traits might affect more complex physiological processes.

In the study presented here, we compared six individuals with four separate and independent EBV-transformations to inquire if EBV-transformation significantly increased the variation within individuals. A better approached would have been to contrast transformed *versus* non-transformed cells. Yet, similar to the LCLs from the 1,000 Human Genome collection, no native (non-transformed) cells are available and thus were not compared; hence we cannot make any conclusions about transacting EBV effects that influence cells regardless of the integration site. Instead, this study addresses whether differences in the integration site or potentially the number of integrations obscured differences among individuals using a nest-analysis of variation: are there significant effects within individuals with separate and independent transformation and does this variation obscure the variation among individuals.

The importance of these data is the ability to address whether the EBV-transformed LCLs found in the 1,000 Human Genome collections can be used to investigate the variation among individual or population; that is, does EBV-transformation obscure potential differences among individuals. From our results, we determined that random viral integration does affect OxPhos metabolism in three of six individuals. State 3 (p=0.0006), E State (p=0.0020), Complex I (p=0.0150), Complex II (p=0.0061), and Complex IV (p=<0.0001) activities all showed significant variation in separate EBV-transformations within an individual. In spite of the effects of viral transformation, significant differences between individuals can still be found in all of these metabolic parameters except for Complex II. It is interesting that Complex II is the only OxPhos enzyme complex with only nuclear and no mitochondrial encoded proteins. The significance of this is unknown but could reflect homeostatic mechanisms between the mitochondrial and nuclear genomes, which could mitigate EBV-transformation effects. Thus, Complex II, which lacks mtDNA-encoded proteins, may be more sensitive to EBV effects. Proton LEAK across the mitochondrial membrane did not appear to have been affected by viral integration (p=0.5634) and also does not appear to be different among individuals (p=0.7074). While the non-productive loss of proton gradient (LEAK) does not have significant variation, it is difficult to address the biological significance. There is likely to be genetic variation in LEAK because both fatty-acid composition and the proteins regulating LEAK have significant heritability [28–30]. Thus, while there are many proteins that affect LEAK, there is little variation in this trait among the six individuals or four EBV-transformations in each individual.

Although, there is a significant effect of EBV-transformation across all individuals, it was only significant in half of our tested individuals (2879, 2881, and 2895). In the other three individuals (2878, 2882, and 2896) there were no significant differences between their independent transformations. Among the EBV-transformations, the effect of transformation tended to be limited (Table 4): significant differences among EBV-transformations were found in only two of the four replicate transformations for two individuals (2879 and 2881) and in several of the replicates in one individual (2995). The weekly order that EBV-transformations were measured and significantly affected one or more OxPhos parameter were similar for two individuals, where the first transformation tested had the lowest OxPhos values. In the third individual, the rank order was different (Table 4): in general the first EBV-transformed cell line had higher OxPhos rates, and the fourth EBV-transformed had the lowest OxPhos rates. The observations that: 1) in only two of the six individuals was the first transformation significantly different, 2) among these two individuals the difference involved different OxPhos parameters and 3) that this trend is not visible in the other four individuals, indicates that order *per se* is not important. None of these tests were corrected for multiple testing to avoid type II error— disregarding an effect when there is one. Using a conservative Bonferroni’s multiple test correction for six statistical tests (i.e., the six OxPhos parameters) in the nested ANOVAs would not alter our results except for E-State, where the p-value difference among individuals is greater than 0.008. For the transformation there are 36 tests (6 individual ANOVAs for 6 parameters) making the critical p-value = 0.0014. With this conservative p-value, EBV-transformation is only significant in one individual (2995) and only for three OxPhos parameters (State 3, E-State and Complex II). Thus, while there are significant EBV-transformation effects across all individuals and transformations (Table 3), few of the independent transformations are significant with conservative multiple test correction. While this multiple correction avoids type I statistical errors (finding a significant difference where none exist), it inflates type II error (disregarding an effect when there is one). For this study, not making a type II error is more important because our concern is that EBV-transformations significantly alters complex physiological phenotypes. Thus, the uncorrected statistical tests are most relevant.

The data presented here suggest that EBV-transformation can affect OxPhos metabolism but only in some OxPhos traits for some individuals. Further, this effect is not large enough to obscure the differences among individuals. Overall, the insight gained from this study provides greater understanding of how EBV integration into the genome of LCLs can influence phenotypes such as the metabolic activity of the OxPhos pathway. It also enhances our knowledge of OxPhos metabolic variation in LCLs, which will be essential for future research in this area of study. Most importantly, these findings indicate that we can effectively study OxPhos metabolism as a phenotype among different EBV-transformed LCLs and compare OxPhos to mRNA expression, nuclear genome sequence variation, and mitochondrial genome sequence variation. These types of studies would enhance our understanding of the genetic variation found in the 1,000 Human Genome collection.

## Data Availability

Raw, and normalize (rates/[DNA]) for all Oxphos measurements for all individuals will be made available in Dyrad. XXYY.

## Acknowledgements

The author would like to thank Dr. Marjorie Oleksiak, and Dr. Danielle McDonald for their guidance and support. This research was funded by the University of Miami Provost Award to D. L. Crawford and grant from NSF IOS 1556396.

## Author contributions

Conceptualization: DLC; Data acquisition: RNW; Statistical analyses: DLC and RNW; Funding acquisition: DLC; Figures: DLC and RNW; Writing original draft: RNW; Writing, review, and editing: DLC and RNW.

